# Explainable machine learning models for glioma subtype classification and survival prediction

**DOI:** 10.1101/2025.06.24.661085

**Authors:** Olga Vershinina, Victoria Turubanova, Mikhail Krivonosov, Arseniy Trukhanov, Mikhail Ivanchenko

## Abstract

Gliomas are complex and heterogeneous brain tumors characterized by an unfavorable clinical course and a fatal prognosis, which can be improved by an early determination of tumor kind. Here, we develop explainable machine learning (ML) models for classifying three major glioma subtypes (astrocytoma, oligodendroglioma, and glioblastoma) and predicting survival rates based on RNA-seq data. Thirteen key genes (*TERT, NOX4, MMP9, TRIM67, ZDHHC18, HDAC1, TUBB6, ADM, NOG, CHEK2, KCNJ11, KCNIP2*, and *VEGFA*) proved to be closely associated with glioma subtypes as well as survival. The Support Vector Machine (SVM) turned out to be the optimal classification model with the balanced accuracy of 0.816 and the area under the receiver operating characteristic curve (AUC) of 0.896 for the test datasets. The Case-Control Cox regression model (CoxCC) proved best for predicting survival with the Harrell’s C-index of 0.809 and 0.8 for the test datasets. Using SHapley Additive exPlanations (SHAP) we reveal the gene expression influence on the outputs of both models, thus enhancing the transparency of the prediction generation process. The results indicate that the developed models could serve as a valuable practical tool for clinicians, assisting them in diagnosing and determining optimal treatment strategies for patients with glioma.

**Simple Summary:** Distinguishing glioma subtypes and assessing patient survival is a non-trivial task due to the high heterogeneity of these brain tumors. Accurate diagnosis is a critical step in developing treatment tactics. In this study, using publicly available RNA sequencing data, we identified a set of key genes and built explainable AI models to classify the major glioma subtypes (astrocytoma, oligodendroglioma, and glioblastoma) and predict patient survival. Experiments evaluating the models demonstrated their ability to generate highly accurate predictions. At the same time, the explainable artificial intelligence approach allowed us to identify relationships between the expression levels of the selected genes and the predictions of the models. Taken together, the obtained results indicate the potential of our predictive models for glioma diagnosis.

## 1. Introduction

Gliomas constitute a group of primary tumors of the central nervous system (CNS) that are widespread in the human population and represent approximately 24% of all primary brain and CNS tumors, as well as 80.9% of malignant tumors [1]. The latest classification by the World Health Organization (WHO) of CNS tumors [2] assigns diffuse gliomas of adult type according to their molecular and genetic characteristics into three main subtypes: astrocytomas (IDH-mutant; II-IV grades), oligodendrogliomas (IDH-mutant and 1p/19q-codeleted; II and III grades), and glioblastomas (IDH-wildtype; IV grade). The typical treatment for glioma involves surgical removal of the tumor, followed by a combination of chemotherapy and radiation therapy. Existing studies show significant differences in overall survival (OS) and response to treatment among molecular subtypes of glioma [3–7].

An accurate determination of the glioma subtype is crucial to personalizing the treatment strategy. However, the ambiguity of histomorphological analysis and the high heterogeneity of the tumors complicate this task [8]. For example, there are well-documented cases of disagreement between different specialists on the histological diagnosis of glioma [9]. In addition, the mutational status may vary in different parts of tumor [10,11], and between patients with the same tumor subtype [12,13]. Therefore, additional strategies and approaches are required for the identification of new biological markers and therapeutic targets, as well as for clinical diagnostics.

Recent advances in high-throughput sequencing have revolutionized the study of molecular changes at the genome-wide level. Sequencing technologies have shown their high performance in diagnosing and classifying tumors, leading to the inclusion of molecular features in the WHO classification in addition to established tumor characterization approaches [2]. At the same time, transcriptomic studies of gliomas often face challenges in linking clinical outcomes. Molecular subtypes of gliomas do not always clearly correlate with response to therapy. In addition, the tumor transcriptome is unstable and changes under the influence of the cell cycle, hypoxia, and therapy. Although informative, analysis of RNA sequencing data requires careful consideration of the biological complexity of gliomas, technical artifacts, and methods capable of analyzing large datasets [14]. To this end, both statistical methods and machine learning (ML) algorithms, including artificial intelligence (AI), are promising tools to analyze massive sequencing data, in particular for various classification and regression tasks.

ML models based on gene expression data have been developed to classify glioblastomas as classical, mesenchymal, and proneural types [15]; to discriminate glioblastoma, diffuse astrocytoma, and anaplastic astrocytoma [16]; to classify and grade astrocytoma, oligodendroglioma, and oligoastrocytoma [17], and some others. In [18], multi-omics data including mRNA expression and DNA methylation were used to identify biomarkers and classify three subtypes of glioma (astrocytoma, oligodendroglioma, and glioblastoma). A similar combination of RNA sequencing and DNA methylation data was used to develop deep learning classifiers for lower-grade gliomas and glioblastoma subtypes [19]. There are also many studies that predict the survival of patients with glioma. This includes models to predict OS in patients with glioblastoma [20], lower-grade glioma [21] or glioma of any grade of malignancy [22].

Often, ML models have complicated and “black-box” architectures that impede the interpretation and explanation of their predictions, affecting end-user trust and application at bedside [23]. Developing eXplainable AI (XAI) systems [24] is particularly necessary in healthcare because such models make critical predictions that can significantly impact treatment and disease outcomes [25]. However, explainable AI models for classification of gliomas refer mainly to MRI image analysis. These include studies on classifying gliomas into two classes (lowand high-grade) [26,27], classifying astrocytoma grades [28], predicting OS in patients with glioblastoma [29], and a few others. It should be noted that many of the cited studies use outdated nomenclatures. According to the 2021 revision, low- and high-grade gliomas are no longer recognized as general categories, which can cause inconsistencies in classification results.

In this work, we develop an explainable ML model based on SHapley Additive exPlanations (SHAP) that classifies three main glioma subtypes — astrocytoma, oligodendroglioma and glioblastoma — based on RNA-seq data. The analysis of the obtained classification model produces an informative set of biological markers (genes) associated with these subtypes. Furthermore, we identify that this set of genes is also associated with patient survival and train the second explainable model to predict OS in patients with glioma, universal with respect to subtype. Gene expression patterns that contribute to the AI classification of glioma subtypes and the prediction of patient survival are described and analyzed.

## 2. Materials and Methods

### 2.1. Data Collection

The bulk RNA-seq data of glioma patients were obtained from two independent data portals. Datasets mRNAseq_693 and mRNAseq_325 were retrieved from the the Chinese Glioma Genome Atlas (CGGA) [30] (http://www.cgga.org.cn/). Datasets TCGA-LGG (low grade glioma) and TCGA-GBM (glioblastoma multiforme) were sourced from The Cancer Genome Atlas (TCGA) [31] (https://portal.gdc.cancer.gov/ and https://www.cbioportal.org/). For each dataset, STAR count files were downloade with the number of mapped RNA-seq reads per gene, as well as the corresponding clinical information. Primary tumor samples were collected for three subtypes of glioma: astrocytoma (class 0), oligodendroglioma (class 1) and glioblastoma (class 2). We used histological classifications following the original clinical data: the “Histology” column for CGGA and the “primary_diagnosis” column for TCGA. Since CGGA data contain all three studied glioma subtypes in a single unified batch, mRNAseq_693 dataset was selected as a training set and mRNAseq_325 dataset was used for internal validation and testing of the models. For additional external tests, the dataset obtained by combining TCGA-LGG and TCGA-GBM data was used.

### 2.2. Data Preprocessing

All datasets were checked for missing values and duplicate samples. The genes identified as protein-coding based on genome annotations GENCODE v19 for CGGA data and GENCODE v36 for TCGA data and common across the datasets were selected for further analysis. Genes that had zero expression in all training samples or exhibited no variation within the training set were excluded from the data. The resulting number of genes after filtering was 17588. Since RNA-seq data have specific properties such as extreme values and mean-variance dependence (heteroscedasticity), all samples were logarithmic transformed *log*_2_(*counts* + 1) to make the data more suitable for ML algorithms. To eliminate technical variations and batch effects between datasets, batch correction was performed using the *ComBat* function [32] from the *sva* R package v.3.50.0 [33], where the mRNAseq_693 dataset (training set) was chosen as the reference batch. Finally, z-score normalization (standardization) to zero mean and unit standard deviation was applied to the training data. The validation and test sets were normalized using the means and standard deviations of the training set. Special normalization methods for RNA-seq data, such as FPKM or TPM, were attempted but not finally used, as they did not improve the quality of predictive models in our case. Interestingly, similar results were observed in [34], comparing machine learning models built on normalized and raw data in the tumor diagnosis task.

### 2.3. Feature Selection

To reduce the dimensionality of the input data space, filter-based feature selection methods were used, which determine the importance of input features by evaluating their relationship to the target variable. Specifically, the association of gene expression with the class label was calculated using the Mutual Information (MI) [35] and Tuned ReliefF (TuRF) [36] methods. To compute MI, we used the *mutual_info_classif* function from the *scikit-learn* Python package v.1.5.1 [37]. The function is based on the estimation of the entropy using distances to the nearest neighbors. To calculate feature importance scores using the TuRF method, we trained the *TuRF* model from the *scikit-rebate* Python package v.0.62 [38]. This model recursively identifies the most important features by examining differences between nearest neighbors. The score increases when the feature values differ between pairs of nearest neighbors belonging to different classes. Feature selection was carried out on the training set, with the number of nearest neighbors kept at the default value for both methods. The features were ranked according to their importance value to select the most significant ones.

### 2.4. ML Models for Classification

Various ML models, both simple and advanced, were evaluated to classify glioma samples into three classes. The k-Nearest Neighbors (kNN) algorithm [39] assigns objects to the class of the majority of their k-nearest neighbors in the training dataset, based on distance metrics such as the Euclidean distance. The Support Vector Machine (SVM) [40] constructs an optimal hyperplane that maximizes the margin between classes, using an “one-vs-rest” strategy for multiclass classification. Decision tree ensembles based on the idea of bagging, including Random Forest (RF) [41] and Extremely Randomized Trees (ERT) [42], train independent decision trees on random bootstrap samples and average their predictions. Decision trees can be combined into an ensemble by boosting, which involves sequentially training base estimators to correct prior errors. Notable Gradient Boosted Decision Trees (GBDT) models include eXtreme Gradient Boosting (XGBoost) [43], Light Gradient Boosting Machine (LightGBM) [44], and Categorical Boosting (CatBoost) [45]. Deep Neural Networks (DNNs) are complex AI systems composed of interconnected layers of nodes, inspired by the structure of the biological neural network. We examined lightweight DNN architectures such as the Attentive Interpretable Tabular Learning Neural Network (TabNet) [46] and the Gated Adaptive Network for Deep Automated Learning of Features (GANDALF) [47].

Software implementations of the models were sourced from the following Python packages: *scikit-learn* v.1.5.1 [37] (kNN, SVM, RF, and ERT), *xgboost* v.2.0.3 [43] (XGBoost), *lightgbm* v.4.3.0 [44] (LightGBM), *catboost* v.1.2.5 [45] (CatBoost), and *pytorch_tabular* v.1.1.0 [48] (TabNet and GANDALF).

To evaluate the performance of classification models on imbalanced data, balanced accuracy (BA) [49] served as the primary metric. The Area Under the Receiver Operating Characteristic (ROC) curve (AUC) was used as an additional performance metric. The LogLoss metric, which measures the divergence between predicted probabilities and actual classes, was used to gauge the calibration of the models. Higher values of the BA and AUC, along with a lower LogLoss, indicate a better performing model. All metrics were calculated using the *scikit-learn* Python package v.1.5.1 [37].

### 2.5. ML Models for Survival Prediction

Establishing an association between survival times and predictor variables, as well as estimating mortality risk scores at given time points, often relies on the Cox Proportional Hazards (CoxPH) regression model [50]. Since Cox regression depends on a linear combination of covariates, it can struggle to capture the nonlinear effects of risk factors on survival, prompting the increasing adoption of more advanced techniques in survival analysis. Random Survival Forest (RSF) [51] and Extra Survival Trees (EST) are ensemble methods that combine multiple survival trees constructed using bootstrap samples and operate similarly to RF and ERT but are adapted to censored survival data. There exists a generalized version of the gradient boosting model known as XGBoost Survival Embeddings (XGBSE)[52]. In this approach, an ensemble of decision trees performs feature transformations (embeddings) on the input data, allowing subsequent training of various models such as logistic regression (XGBSEDebiasedBCE, abbreviated as XGBSE-DBCE) or kNN (XGBSEKaplanNeighbors, abbreviated as XGBSE-KN). DNN architectures are also used to predict survival. Neural networks such as the Cox proportional hazards deep neural network (DeepSurv) [53] and the Case-Control Cox regression model (CoxCC) [54] are nonlinear extensions of Cox regression and differ in their loss functions, which are minimized. The Piecewise Constant Hazard model (PC-Hazard) [55] assumes that the continuous-time hazard function is constant within predefined intervals. The Neural Multi-Task Logistic Regression model (N-MTLR) [56] is a discrete-time approach that models the hazard function by combining multiple locally dependent logistic regression models, utilizing nonlinear data transformations from a multilayer perceptron.

Software implementations of the models were sourced from the following Python packages: *scikit-survival* v.0.23.0 [57] (CoxPH, RSF, and EST), *xgbse* v.0.3.1 [52] (XGBSE-DBCE and XGBSE-KN), and *pycox* v.0.3.0 [54] (DeepSurv, CoxCC, PC-Hazard, and N-MTLR).

The Harrell’s concordance index (C-index) [58], which measures the rank correlation between predicted and observed outcomes, was used as the primary metric to evaluate the performance of survival prediction models. We also analyzed the time-dependent ROC curve [59], a modified version of the traditional ROC curve for time-to-event data, and the associated AUC values. Furthermore, we evaluated model calibration using the Brier score [60], calculating the Integrated Brier Score (IBS) for a comprehensive assessment at all time points. Higher values of the C-index and AUC, along with a lower IBS, indicate a better performance model. The C-index and IBS were computed using the respective Python packages for the models. Time-dependent ROC analysis for all models was performed using the *survivalROC* R package v.1.0.3.1 [59].

### 2.6. Tuning and Training ML Models

For each ML model, hyperparameter tuning was performed on the training dataset (CGGA mRNAseq_693), using a stratified 5-fold cross-validation approach. For the classification task, stratification was based on the class label, while for the survival prediction task, stratification was based on survival status. To identify the most effective configuration of hyperparameters, we utilized the multivariate Tree-structured Parzen Estimator (TPE) algorithm with the group decomposed search space [61,62] (the software implementation was sourced from the *optuna* Python package v.3.6.1 [63]). Details concerning the tunable hyperparameters for each model–including their descriptions and corresponding search distributions–are presented in Supplementary Table S1. The total number of hyperparameter optimization trials for each experiment was set to 500. The combination of hyperparameters that provided the highest cross-validation metric (BA for the classification problem, C-index for survival prediction) was considered optimal.

The CGGA mRNAseq_325 dataset was randomly divided equally into validation and internal test sets using class stratification. The validation dataset was used to monitor the training of the GBDT and DNN models during the early stopping process: the maximum number of training rounds was set to 100, and the number of early stopping rounds was set to 10. Training was terminated if the evaluation metric in the validation dataset did not improve for 10 consecutive rounds. By introducing a separate validation dataset that is distinct from both the training and final test datasets, early stopping can be efficiently implemented, preventing overfitting, and ensuring generalization. Regularly evaluating the behavior of models on previously unseen data helps determine when further training ceases to improve prediction accuracy, thereby saving computational resources and maintaining robustness against errors.

For training DNN models, the Adam optimization algorithm [64] was used and the batch size was fixed at 32. For the TabNet and GANDALF models, the automatic selection option of the optimal learning rate was enabled, while for the DeepSurv, CoxCC, PC-Hazard and N-MTLR models, the learning rate was determined by hyperparameter tuning.

The trained models with optimized hyperparameters were subsequently evaluated on an independent internal test dataset and an external test dataset to identify the model that exhibited superior performance.

### 2.7. eXplainable Artificial Intelligence

The SHAP (SHapley Additive exPlanations) explainability method is a game theory-based approach that quantifies the impact of features on predictions by assigning Shapley values to them [65]. The SHAP method is particularly useful for interpreting complex “black-box” ML models. SHAP values are derived by averaging the contributions of a feature to the model’s predictions, considering all possible permutations of other features within the model. SHAP values show how changing a specific feature value for a given data point affects the model’s base prediction (which is usually the average prediction over the training dataset). Using the SHAP method, both global and local explainability can be obtained. Global explainability refers to the analysis of aggregate contributions from features in a dataset. This helps understanding the general patterns of influence of various features on the model predictions as a whole. Local explainability focuses on explaining the contribution of features to individual predictions in order to provide an understanding of the reasons behind the specific decisions made by the model. To calculate the SHAP values, the *Explainer* function from the *shap* Python package v.0.45.1 was used with the training dataset as a background.

### 2.8. Functional Enrichment Analysis

Gene Ontology (GO) enrichment analysis of the gene set was performed using the online tool g:Profiler [66] (https://biit.cs.ut.ee/gprofiler/gost). GO terms were considered statistically significantly enriched if the adjusted p-value after Benjamini–Hochberg (BH) correction was less than 0.001. In addition, redundant terms were eliminated using the greedy search strategy implemented in g:Profiler. To evaluate the enrichment level of significant GO terms, single-sample gene set enrichment analysis (ssGSEA) was carried out using the *GSVA* R package v.1.50.5. This method assess the enrichment of gene sets included in specific GO terms in various samples and, consequently, across distinct tumor subtypes. The Mann–Whitney U test with p-values adjusted by the BH procedure was used to compare the ssGSEA scores between the three subtypes of glioma. For post hoc pairwise comparisons, the Dunn test with adjusted p-values was used.

### 2.9. Statistical Analysis

Statistical analysis was performed using the *scipy* v.1.13.1 and *lifelines* v.0.28.0 [67] Python packages. Comparison of clinical characteristics between datasets used the MannWhitney U test for continuous variables and the *χ*^2^ test for categorical variables. An adjusted p-value (after Benjamini–Hochberg correction) less than 0.05 was considered statistically significant. A univariate Cox regression analysis with p-value adjustment tested the association between gene expression levels and overall survival (OS) of the patients. Kaplan-Meier survival curves and the log-rank test were used to compare differences in OS between the two mortality risk groups. The low and high risk groups were defined based on the median risk score from the training dataset.

## 3. Results

### 3.1. Patient Characteristics

ML models for the diagnosis of glioma were based on publicly accessible RNA sequencing data from primary glioma samples that were acquired from the CGGA and TCGA portals. The clinical characteristics of the assembled patient cohorts are summarized in Table 1. No statistically significant differences were found between the training and test datasets from the CGGA. At the same time, most of the clinical variables were significantly different between the training data and the external TCGA test set. In particular, patients from the TCGA tended to be older and had a shorter median overall survival.

**Table 1.** Clinical characteristics of glioma patient cohorts included in the study.

| Characteristic | CGGA,<br>mRNAseq_693<br>(training set,<br>n = 398) | CGGA,<br>mRNAseq_325<br>(validation/test set,<br>n = 229) | TCGA,<br>TCGA-LGG and<br>TCGA-GBM<br>(external test set,<br>n = 536) | p adjusted<br>(validation/test set<br>vs. training set) | p adjusted<br>(external test set<br>vs. training set) |
| --- | --- | --- | --- | --- | --- |
| **Histology** |  |  |  | 0.590 | < 0.001 |
| Astrocytoma | 167 | 84 | 193 |  |  |
| Oligodendroglioma | 91 | 60 | 190 |  |  |
| Glioblastoma | 140 | 85 | 153 |  |  |
| **Grade** |  |  |  | 0.058 | 0.213 |
| WHO II (G2) | 130 | 94 | 154 |  |  |
| WHO III (G3) | 128 | 50 | 187 |  |  |
| WHO IV (G4) | 140 | 85 | 153 |  |  |
| Unknown | 0 | 0 | 42 |  |  |
| **IDH mutation**<br>**status** |  |  |  | 0.704 | 0.125 |
| Wildtype | 175 | 112 | 216 |  |  |
| Mutant | 200 | 116 | 313 |  |  |
| Unknown | 23 | 1 | 7 |  |  |
| **MGMT promoter**<br>**methylation** |  |  |  | 0.058 | < 0.001 |
| Methylated | 185 | 99 | 365 |  |  |
| Unmethylated | 136 | 116 | 140 |  |  |
| Unknown | 77 | 14 | 31 |  |  |
| **Age, years** |  |  |  | 0.823 | < 0.001 |
| Range | 11–76 | 10–79 | 17–89 |  |  |
| Median | 43 | 43 | 48 |  |  |
| **Gender** |  |  |  | 0.590 | 0.926 |
| Male | 233 | 142 | 311 |  |  |
| Female | 165 | 87 | 225 |  |  |
| **Survival status** |  |  |  | 0.115 | < 0.001 |
| Alive | 179 | 85 | 313 |  |  |
| Dead | 204 | 139 | 221 |  |  |
| Unknown | 15 | 5 | 2 |  |  |
| **OS, days** |  |  |  | 0.315 | < 0.001 |
| Range | 27–4725 | 19–4809 | 1–6423 |  |  |
| Median | 1283 | 1118 | 544 |  |  |

WHO: World Health Organization, IDH: isocitrate dehydrogenase, OS: overall survival.

### 3.2. Selection of Features Associated With Glioma Subtypes

RNA-seq data are characterized by being high-dimensional and low-sample-size (HDLSS), leading to increased computational costs when training ML models. In addition, high-dimensional data may contain many redundant and irrelevant features (genes), which degrade predictive performance. To overcome this challenge, we applied two methods to reduce the dimensionality of the feature space. Mutual Information (MI) criterion and the TuRF algorithm provided feature importance scores by quantifying the relationship between feature expression values and class labels (class 0—astrocytoma, class 1—oligodendroglioma and class 2—glioblastoma). In each method, the features were sorted in descending order of importance score, and 1% of the top features (176 features) was selected. In order to improve the reliability and relevance of the shortlisted features we kept only 84 of them found in the intersection (see Supplementary Table S2).

### 3.3. Classifier for Glioma Subtypes

We applied nine ML models to classify the subtypes of glioma based on the z-score of the normalized expression of 84 selected genes. For each model, hyperparameter optimization was performed, followed by training the models with their optimal combinations. The performance of the models was evaluated using balanced accuracy (BA) metrics, which was estimated by stratified 5-fold cross-validation and on internal and external test datasets (Table 2).

**Table 2.** Results of glioma subtype prediction based on 84 genes. Models are ranked based on their balanced accuracy computed on the test dataset in descending order.

| Model | BA, train | BA, CV | BA, test | BA, external test |
| --- | --- | --- | --- | --- |
| GANDALF | 0.793 | **0.824** | **0.808** | 0.785 |
| kNN | **0.802** | **0.805** | **0.805** | 0.794 |
| SVM | **0.843** | **0.823** | **0.805** | **0.821** |
| LightGBM | **0.852** | **0.833** | 0.778 | 0.781 |
| TabNet | **0.830** | **0.826** | 0.777 | 0.784 |
| XGBoost | **0.843** | **0.826** | 0.775 | 0.775 |
| RF | **0.934** | **0.821** | 0.772 | 0.780 |
| ERT | **0.917** | **0.827** | 0.767 | 0.788 |
| CatBoost | 0.783 | **0.842** | 0.714 | 0.728 |

BA: balanced accuracy, CV: cross-validation. Values greater than 0.8 are highlighted in bold.

All models showed high cross-validation accuracy (above 0.8), and only three models (GANDALF, kNN, and SVM) demonstrated accuracy above 0.8 on an internal test dataset that was not included in the training process. At the same time, only the SVM model achieved a performance greater than 0.8 on the external test dataset, demonstrating a superior level of generalization. Moreover, the SVM model had one of the highest AUC values among the others (see Supplementary Table S3). Furthermore, the SVM model demonstrated one of the lowest LogLoss scores, indicating good calibration and strong agreement between its predicted probabilities and the actual class labels.

To reduce the number of variables in the SVM model and improve its applicability, an additional round of feature selection was conducted. We sorted the 84 features in descending order according to their importance for global explainability, as determined by SHAP values. The global importance of the features was estimated by averaging the absolute SHAP values computed for the training dataset. We then considered a series of feature subsets, including 1, 2, …, 84 the most important features. For each subset, hyperparameter tuning, training, and testing of the SVM model were conducted. We achieved the highest BA on the test dataset, *BA*_*test*_ = 0.816, when the top 13 genes (*TERT, NOX4, MMP9, TRIM67, ZDHHC18, HDAC1, TUBB6, ADM, NOG, CHEK2, KCNJ11, KCNIP2*, and *VEGFA*) were used to train the SVM classifier with polynomial kernel (Figure 1A). The classification accuracy calculated for the training dataset using cross-validation and for the external test dataset was also high: *BA*_*CV*_ = 0.837 and *BA*_*external test*_ = 0.816, respectively. Therefore, the 13-gene SVM model was selected as the optimal model for the classification of gliomas.

**Figure 1.**
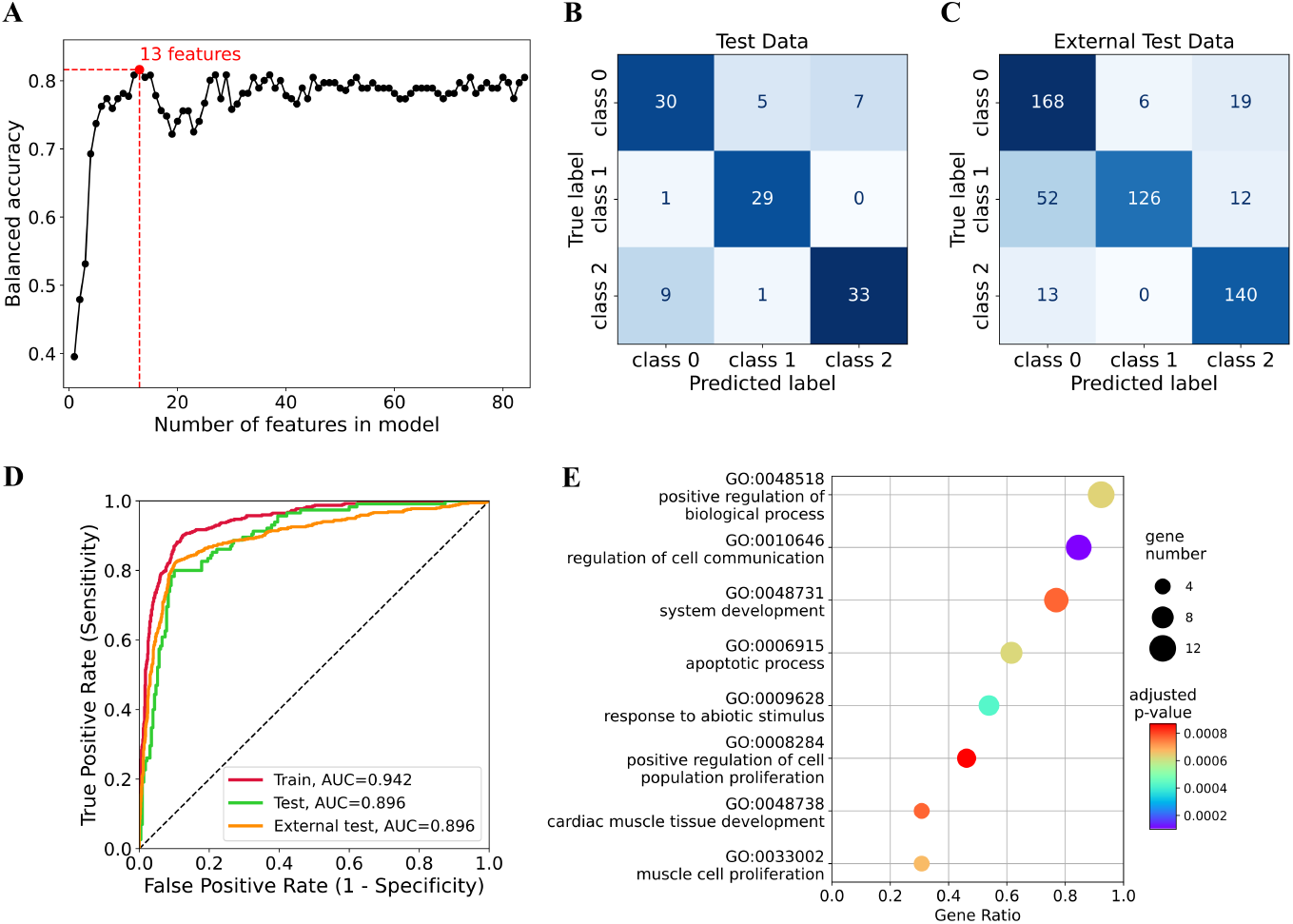
Establishment and validation of the optimal classification SVM model for three glioma subtypes. (**A**) Dependence of the balanced accuracy calculated on the internal test dataset on the number of features in the model. Dotted lines correspond to the optimal small model with a balanced accuracy of 0.816 and 13 features. (**B**) Confusion matrix of the optimal 13-gene model for the internal test dataset. (**C**) Confusion matrix of the optimal 13-gene model for the external test dataset.(**D**) ROC curves for the optimal 13-gene model. ROC curves were generated using training, internal test and external test datasets. (**E**) GO enrichment analysis of 13 selected genes. Class 0: astrocytoma, class 1: oligodendroglioma, class 2: glioblastoma.

The confusion matrices for the resulting classifier model are shown in Figure 1B,C. In both test datasets, the tumor subtypes of most patients are correctly predicted, as indicated by the high values of the diagonal elements. We observed misclassifications between astrocytomas and glioblastomas. In addition, some oligodendrogliomas were misclassified as astrocytomas by our model on the external test dataset. Transcriptomic data from a single glioma subtype may show significant variation due to intratumor heterogeneity resulting from clonal evolution, microenvironment (hypoxia, immune infiltration), tumor region, and epigenetic modifications. Polymorphisms in the regulatory regions of genes can also influence expression. It should be taken into account that the studied databases rely on the nomenclature accepted before 2021. IDH-mutated diffuse astrocytic tumors were then classified into 3 different tumor types (diffuse astrocytoma, anaplastic astrocytoma, and glioblastoma) based on histological parameters. However, in the current classification, all diffuse astrocytic tumors with the IDH mutation are considered one type (astrocytoma, IDH mutation) and are classified as grades 2, 3 or 4 on the WHO CNS scale. The model may classify some astrocytomas as glioblastomas since they are similar in molecular profile, aligning with the 2021 WHO classification. The rise in classification errors observed in the external test dataset could be due to inconsistencies in the class annotations compared to the training data.

The relationship between sensitivity and specificity of the classification model is presented in Figure 1D by the ROC curves. The AUC values for both test datasets were high (*AUC*_*test*_ = *AUC*_*external test*_ = 0.896), indicating that the 13-gene SVM model can classify glioma subtypes with high specificity and sensitivity, and has potential at the bedside.

### 3.4 Functional Analysis of Prognostic Genes

The GO enrichment analysis (Figure 1E) performed using online tool g:Profiler revealed that in the “biological processes” category, the 13 prognostic genes were significantly enriched in positive regulation of biological process (GO:0048518), regulation of cell communication (GO:0010646), system development (GO:0048731), apoptotic process (GO:0006915), response to abiotic stimulus (GO:0009628), positive regulation of cell population proliferation (GO:0008284), cardiac muscle tissue development (GO:0048738), and muscle cell proliferation (GO:0033002). We employed the ssGSEA method to quantify the enrichment levels of these significant biological processes in glioma samples. For this purpose, subsets of 13 genes included in the corresponding GO terms were extracted from the g:Profiler output. The terms GO:0010646, GO:0006915, GO:0009628, and GO:0008284 showed significant enrichment in glioblastomas, while the terms GO:0048518, GO:0048731, GO:0048738, and GO:0033002 were enriched in oligodendrogliomas (Supplementary Figure S1). No significant enrichment was detected in the astrocytomas. The findings suggest different molecular mechanisms driving each subtype and require further investigation.

### 3.5. Model for Survival Prediction

We then examined the relationship between the 13 identified genes and overall survival (OS) rates in patients. The maximum follow-up time in the training dataset was 4725 days (12.9 years). Patients from both test datasets whose survival times were outside the ranges of the training dataset were excluded. According to the results of the univariate Cox regression analysis, all genes demonstrated significant associations with OS in both the training and test cohorts, cf. Supplementary Table S4. Hazard ratios (HR) less than 1 for covariates *TRIM67, NOG, KCNJ11*, and *KCNIP2* indicated a positive association with OS, while HR greater than 1 for the remaining covariates corresponded to a negative association with OS.

Based on these observations, we used 13 previously isolated genes to construct a survival predictive model in a mixed cohort. We tuned hyperparameters and trained nine ML models, encompassing both traditional multivariate Cox regression and more sophisticated algorithms such as survival forests and neural networks. The evaluation of the performance of the models is presented in Table 3 and Supplementary Table S5.

**Table 3.** Results of glioma patient survival prediction based on 13 genes. Models are ranked in descending order according to their C-index scores computed on the test dataset.

| Model | C-index, train | C-index, CV | C-index, test | C-index, external test |
| --- | --- | --- | --- | --- |
| RSF | **0.851** | 0.781 | **0.815** | **0.816** |
| PCHazard | **0.811** | 0.787 | **0.814** | 0.783 |
| EST | **0.810** | 0.779 | **0.813** | **0.815** |
| CoxPH | 0.783 | 0.765 | **0.812** | **0.803** |
| CoxCC | **0.802** | 0.790 | **0.809** | **0.800** |
| XGBSE-DBCE | **0.919** | 0.778 | **0.804** | 0.799 |
| XGBSE-KN | **0.858** | 0.784 | **0.800** | **0.812** |
| N-MTLR | **0.808** | 0.787 | 0.796 | 0.784 |
| DeepSurv | **0.800** | 0.787 | 0.787 | 0.786 |

CV: cross-validation. Values greater than 0.8 are highlighted in bold.

RSF, EST, CoxPH, CoxCC, and XGBSE-KN models consistently achieved high C-index values exceeding 0.8 on both test datasets. However, the models with the highest C-index in the test data (RSF, EST, and CoxPH) also exhibited a high IBS value, suggesting that they are poorly calibrated (see Supplementary Table S5). In result, the CoxCC neural network was selected as the best model based on its high C-index value on the test datasets (*C*-*index*_*test*_ = 0.809, *C*-*index*_*external test*_ = 0.8) and the highest cross-validation score for the metric (*C*-*index*_*CV*_ = 0.790). In addition, CoxCC demonstrated low values of IBS.

The selected model was then used to predict the mortality risk score for each patient. According to the median risk score calculated on the training dataset, patients were divided into low-and high-risk groups. To assess the discriminatory ability of the model, we performed a survival analysis. It showed that OS in the low-risk group was significantly higher compared to the high-risk group (log-rank p-value < 0.05) in both test datasets (Figure 2A,B). This is also confirmed by HR analysis: patients with the low-risk score have a better prognosis in both the internal test set (*HR* (95% *CI*) = 2.92 (2.23 − 3.82)) and the external test set (*HR* (95% *CI*) = 2.8 (2.4 − 3.26)).

**Figure 2.**
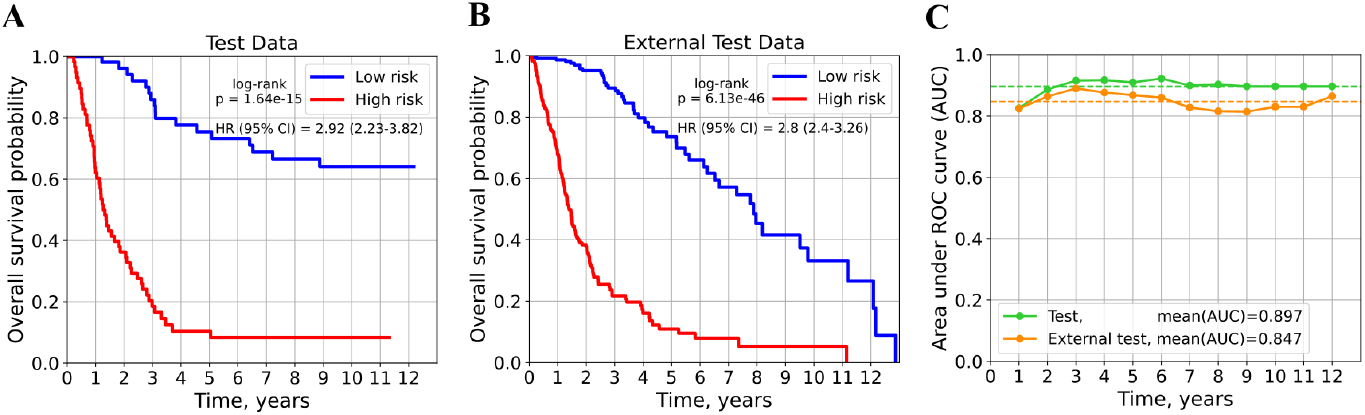
Validation of the CoxCC model for predicting overall survival in patients with glioma. **(A)** Kaplan–Meier survival curves for the low- and high-risk groups in the internal test dataset. **(B)** Kaplan–Meier survival curves for the low- and high-risk groups in the external test dataset. **(C)** Time-dependent ROC analysis of the prognostic model in the test datasets. The dotted lines represent the mean AUC value calculated over 12 years.

To further evaluate the predictive performance of the model, we performed a time-dependent ROC analysis by calculating the areas under the ROC curves to predict risk scores over a period of 1 to 12 years (Figure 2C). This approach allowed us to estimate how accurately the risk score determined by the model can discriminate patients who died by a specified period (e.g., 1, 2, …, 12 years) from those who remained alive. In the internal test dataset, the AUC values were 0.910 and 0.897 to predict 5- and 10-year OS, respectively, and the mean AUC value over 12 years was 0.897. In the external test dataset, the values were slightly lower but remained high: the AUCs for 5- and 10-year predictions were 0.868 and 0.830, respectively, and the mean AUC value was 0.847.

The results obtained demonstrated the reliability and high predictive power of the CoxCC neural network trained on 13 genes in predicting the survival outcomes of patients with glioma.

### 3.6. Explainability of ML models

To understand how the resulting models make predictions, we used a SHAP explainability method to assess the importance of features and their contributions to diagnostics. By examining SHAP values, one can determine which features have the greatest impact on particular outcomes and infer the rationale behind specific predictions. In our study, we illustrate the global and local explainability of our best models using samples from the internal test dataset. To calculate the SHAP values for the glioma subtype classification model, the class probabilities predicted by the SVM model were utilized. In the context of multi-class classification, the SHAP values are calculated for each individual class. For the survival analysis model, the 12-year survival probability predicted by the CoxCC model was used.

The global explainability of the models is illustrated in Figure 3. In the plots, the prognostic genes are ordered by their average contribution to the output of the models. For the classification model of three glioma subtypes (Figure 3A), the contribution of features to the prediction of each class is indicated by different colors. For example, the *NOX4* gene has the greatest influence on classification, and its expression levels are more critical for predicting astrocytomas and glioblastomas than oligodendrogliomas. For the survival prediction model (Figure 3B), gene *NOG* is the most important. It is worth noting that the order of importance of genes for the two models differs significantly. For instance, gene *MMP9* exhibits high importance for classification but low importance for survival prediction, whereas gene *KCNJ11* demonstrates the opposite trend, showing high importance for survival prediction and low importance for classification.

**Figure 3.**
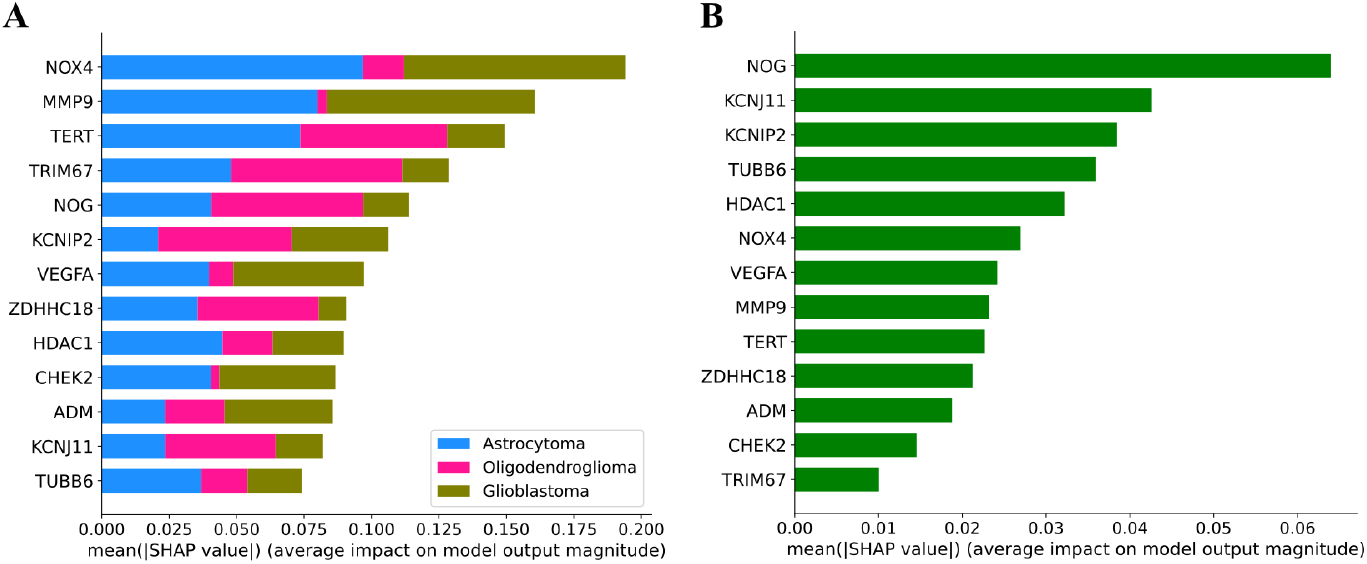
Average impact (the mean absolute SHAP values) of features on model predictions. Features with higher values exert stronger influence. (**A**) Global importance of features in the classification model for three glioma subtypes. For each feature, its contribution to certain classes is indicated by the corresponding color. (**B**) Global importance of features in the model for predicting overall survival of patients with glioma.

Then we constructed scatterplots to visualize the direction of the relationship between the prediction of the model and the expression levels of the most important genes (Figure 4) and the remaining genes (Supplementary Figures S2 and S3).

**Figure 4.**
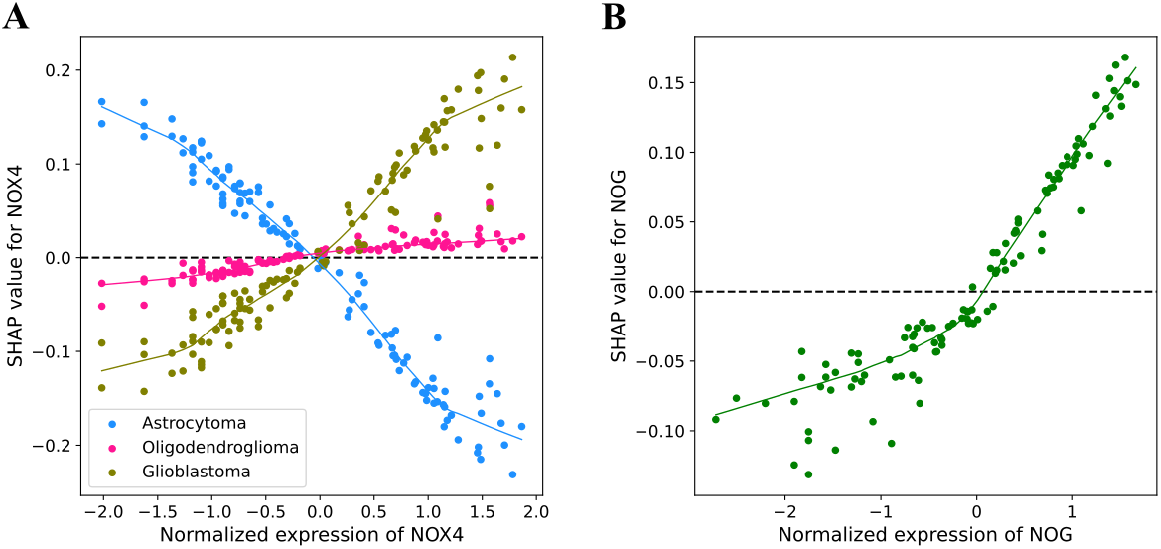
Scatterplots for the SHAP values and expression levels of the most important genes in the models. (**A**) Dependence of SHAP values on the expression of the *NOX4* gene in the classification model for three glioma subtypes. Each patient in the test dataset corresponds to three dots, which indicate the SHAP values associated with the three glioma subtypes. Positive SHAP values for a particular class indicate that the feature value increases the probability of predicting that class, while negative values decrease it. (**B**) Dependence of SHAP values on the expression of the *NOG* gene in the model for predicting overall survival of patients with glioma. Positive SHAP values indicate that the feature value increases overall survival, while negative values decrease it.

In a classification problem, the SHAP value indicated how much the probability of predicting a certain class increases (if the SHAP value is positive) or decreases (if the SHAP value is negative), depending on the observed value of the considered feature. As shown in Figure 4A, high levels of *NOX4* expression increased the probability of predicting glioblastoma, as evidenced by the positive correlation between the SHAP values for the glioblastoma class and the gene expression values. Low *NOX4* expression values increased the probability of predicting astrocytoma, as a negative correlation is observed between the SHAP values for the astrocytoma class and gene expression. SHAP values for the oligodendroglioma class were close to zero, indicating a minimal impact of gene expression on this tumor subtype. Analysis of the SHAP values for the remaining genes (Supplementary Figure S2) revealed that increased expression of *ZDHHC18, HDAC1*, and *TUBB6* enhances the probability of predicting astrocytoma, while elevated expression of *TERT, TRIM67, NOG, KCNIP2*, and *KCNJ11* increases the probability of predicting oligodendroglioma. In addition, higher expression levels of *MMP9, TERT, VEGFA, ZDHHC18, CHEK2*, and *ADM* elevate the probability of predicting glioblastoma.

For the survival prediction model, SHAP values determine the amount of increase (or decrease) in the predicted survival probability due to a given feature value. We observed that as the expression of the *NOG* gene increased, the SHAP values increased monotonically, indicating better patient survival (Figure 4B). Regarding the remaining genes (Supplementary Figure S3), increased expression of *KCNJ11* and *KCNIP2* also associated with increased survival, while increased expression of *TUBB6, HDAC1, NOX4, VEGFA, MMP9, TERT, ZDHHC18, ADM*, and *CHEK2* reduced survival. The relationship between mortality risk and *TRIM67* expression was highly nonlinear.

Using SHAP values, individual model predictions can be explained locally. The SHAP waterfall plots in Figure 5 illustrated the local explainability of our SVM classification model for a correctly classified glioblastoma sample. The graphs demonstrated how the model adjusts its probability prediction for each class based on patient-specific normalized expression values. The genes in the graphs are arranged from bottom to top according to their increasing importance to refining the prognosis. The blue (red) lines correspond to features whose values decrease (increase) the probability of the corresponding class. For the sample considered, the expression levels of 11 of 13 genes increase the probability of the “Glioblastoma” class (Figure 5C), resulting in the prediction of this tumor subtype with a probability of 0.948.

**Figure 5.**
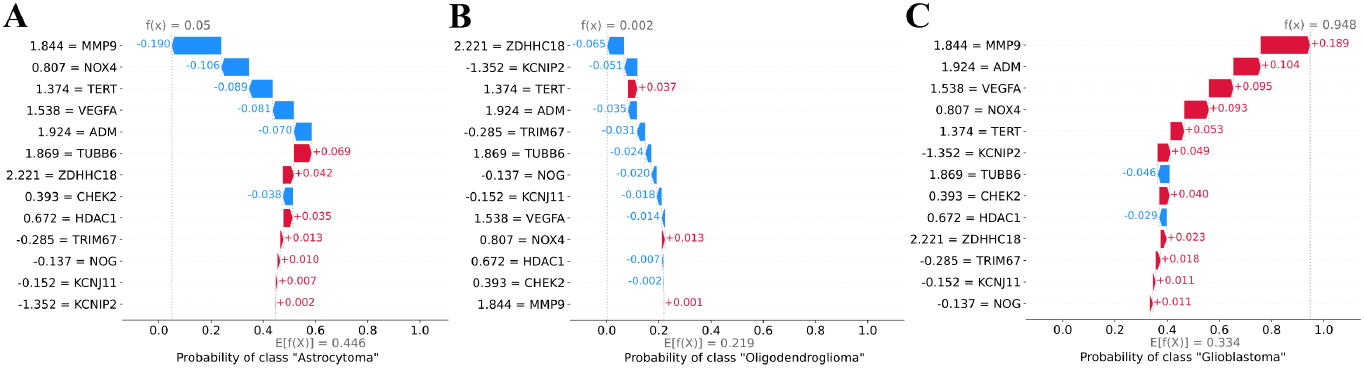
SHAP waterfall plots of an individual sample from the test dataset for a model classifying the glioma subtype. (**A**) Waterfall plot for class “Astrocytoma”. (**B**) Waterfall plot for class “Oligodendroglioma”. (**C**) Waterfall plot for class “Glioblastoma”. The lower part of the graphs displays the average (baseline) probability of predicting the corresponding class in the training dataset, *E*[*f* (*X*)]. From bottom to top, features are presented in order of their increasing contribution to the prediction, with each line indicating how the feature value modifies the prediction probability. Blue lines represent negative SHAP values and show how much the feature value decreases the probability of the class, whereas red bars represent positive SHAP values and show how much the feature value increases the probability of the class. The resulting *f* (*x*) values are the probabilities of the classes predicted by the model, with the highest one determining the class to which the tested sample will be assigned.

Figure 6 displays local explainability graphs of survival predictions generated by the CoxCC model for three distinct patients with glioma. The blue (red) lines correspond to features whose values decrease (increase) the survival probability. For patients who died after 450 and 1133 days, the model predicted low 12-year survival probabilities of 0.019 (Figure 6A) and 0.45 (Figure 6B), respectively. Meanwhile, for the patient who survived just over 12 years (4427 days), normalized expression levels of all genes were indicative of prolonged survival, which led the model to predict a high survival probability of 0.84 (Figure 6C).

**Figure 6.**
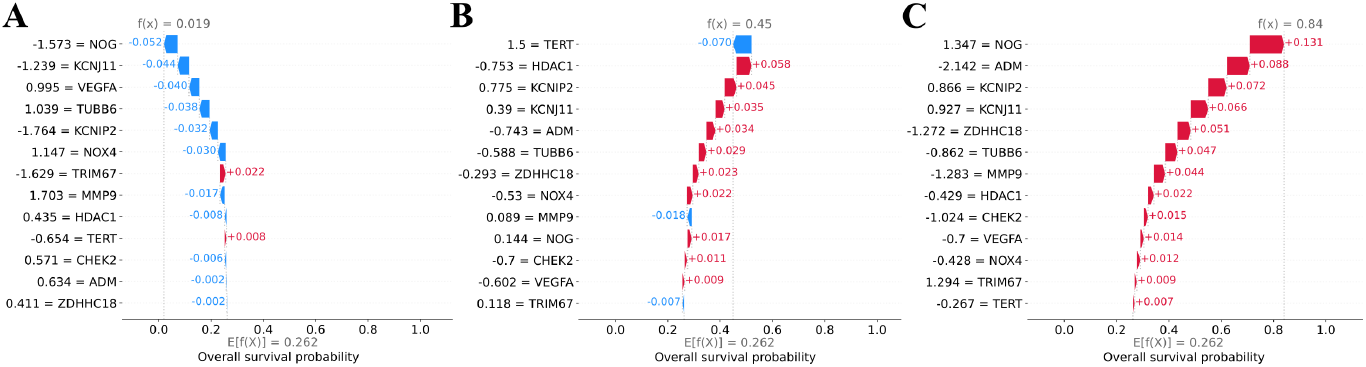
SHAP waterfall plots of individual glioma patients from the test dataset for a model predicting survival. (**A**) Waterfall plot for a patient who died after 450 days. (**B**) Waterfall plot for a patient who died after 1133 days. (**C**) Waterfall plot for a patient who was alive at least 4427 days later. The lower part of the graphs displays the average (baseline) predicted survival probability in the training dataset, *E*[*f* (*X*)]. From bottom to top, features are presented in order of their increasing contribution to the prediction, with each line indicating how the feature value modifies the prediction probability. Blue lines represent negative SHAP values and show how much the feature value decreases the survival probability, whereas red bars represent positive SHAP values and show how much the feature value increases the survival probability. The resulting *f* (*x*) values represent the 12-year survival probabilities predicted by the model.

## 4. Discussion

In this study, we developed the ML model based on the expression of protein-coding genes to classify three subtypes of glioma with varying degrees of severity: astrocytoma, oligodendroglioma, and glioblastoma. Feature selection identified the most important genes associated with subtypes. Among the nine classification algorithms evaluated, the SVM model trained on 13 key genes demonstrated the highest performance with balanced accuracy of 0.816 and AUC of 0.896 in both test datasets. Despite the existence of more advanced ML algorithms, such as gradient boosting and neural networks, the SVM model has also previously shown a high efficiency in classifying gliomas [16,17, 68]. The implementation of SHAP, an explainable AI technique, enabled visualization of the magnitude and direction of the contribution of specific input variables to the model predictions for the cohort in general and individual patients. Some existing explainable classification models focus mainly on categorizing gliomas into two classes based on MRI images [26,27,69]. We found only one study that classifies gliomas into the same three classes as we did and used various libraries to visualize explainability [70]. However, that work relies on gene mutation data, whereas we use RNA-seq data.

In addition, we examined the relationship between the expression levels of the 13 identified genes and patient survival in the mixed cohort. The presence of a statistically significant association of gene expression with survival allowed us to build the ML model to predict the probability of survival. The resulting model based on the CoxCC neural network demonstrated high predictive capability. Thus, the C-index achieved values of 0.809 and 0.800 for the internal and external datasets, respectively. Furthermore, the 5-and 10-year AUC were 0.910 and 0.897 for the internal dataset, and 0.868 and 0.830 for the external dataset. To date, numerous studies have explored prediction glioma patient survival using RNA-seq data, although most rely primarily on the Cox regression model [20–22,71–73]. The advantage of our model lies in its superior predictive quality, outperforming most of the existing models, combined with the global and local explainability for the predictions it generates.

According to the literature, the genes identified exhibit a strong association with glioma subtypes and OS. mRNA expression levels of *NOX4* in glioblastomas (WHO grade IV) were significantly higher compared to astrocytomas (WHO grades II and III) [74], and OS was significantly lower in patients with high *NOX4* expression [75]. *MMP9* expression levels are increased in high-grade gliomas and are overexpressed in glioblastomas [76,77], while lower *MMP9* expression correlates with improved patient survival [76–78]. *KCNIP2* gene expression was elevated in grade II and III gliomas compared to grade IV gliomas, and its high expression was significantly associated with prolonged OS [79]. *KCNJ11* expression was related to the degree of tumor malignancy and was an independent factor that affected the OS, and survival times of patients with high expression of KCNJ11 were longer than those of patients with low expression [80]. Increased *VEGFA* expression is significantly associated with decreased OS in patients with glioblastoma [81]. Interestingly, 4 genes *MMP9, KCNIP2, KCNJ11* and *VEGFA* were identified by the authors of [82] as prognostic markers for glioma-related epilepsy. Increased expression of *NOG*, included in a number of prognostic models, leads to an increase in OS in patients with glioma [83,84]. *TRIM67* is known to be a highly expressed gene in oligodendrogliomas [85]. In work [86], it was mentioned that increased expression of *ZDHHC18* leads to decreased OS in LGG. As glioma grade increased, *ADM* mRNA expression levels increased and this increase was associated with poorer OS [87]. High *HDAC1* expression was significantly associated with the clinical stage of glioma and reduced OS [88]. Increased *TUBB6* gene expression is associated with decreased OS in patients with primary glioblastoma [89].

To our knowledge, there is no evidence of an association between the expression levels of the genes *TERT* and *CHEK2* and glioma subtypes. However, it’s well-established that the *TERT* mutation, commonly seen in glioblastomas and oligodendrogliomas, is informative for the classification of CNS tumors [2]. It is known that *TERT* activity is associated with telomerase function and is often elevated in cancer cells, allowing them to divide indefinitely. No clear correlations were shown between the length of the telomere and the type of glioma, but more malignant tumors with low OS had shorter telomeres than less malignant tumors with longer OS [90,91]. In glioblastomas, median OS is significantly lower than in other subtypes, and this factor could be indirectly related to the transcriptomic activity of the TERT identified in our classifier. Mutations in the *CHEK2* gene have been identified in glioblastomas [92,93] and glial CNS tumors in children [94]. Mutations in the CHEK2 gene may be a risk factor for gliomas, but this link requires further study. More research is needed to clarify the role of this gene in the development of glial tumors.

Functional analysis of 13-gene signature revealed significant associations with tumor progression and cancer-related pathways. There were marked differences in the enrichment of GO terms between the three subtypes of glioma. In glioblastomas, the terms most significantly enriched were regulation of cell communication, apoptotic process, response to abiotic stimulus, and positive regulation of cell population proliferation. Oligodendrogliomas showed enrichment in the following terms: positive regulation of biological processes, system development, cardiac muscle tissue development, and muscle cell proliferation. Unlike glioblastomas and oligodendrogliomas, astrocytomas did not show any significantly enriched GO terms. The existing results corroborate our findings. Multiple studies have highlighted the critical role of intercellular communication in glioblastomas [95–97], which appears to facilitate tumor proliferation and progression. In particular, malignant cells can autonomously stimulate their own growth and division, effectively bypassing normal regulatory mechanisms. The presence of signaling pathways associated with increased proliferative activity in glioblastomas [98,99] probably reflects the inherent aggressiveness of these tumors, which exhibit rapid proliferation and extensive tissue infiltration. These activated intercellular interactions and increased proliferative capacity collectively sustain glioblastoma’s aggressive phenotype and invasive growth patterns. Interestingly, IDH-mutated gliomas demonstrate significant enrichment of the two GO terms: positive regulation of biological processes and system development [100]. These findings suggest distinct molecular mechanisms between the glioma subtypes and have important implications for understanding glioma pathogenesis. Subtype-specific enrichment patterns indicate that the genes identified can serve as potential diagnostic biomarkers. However, additional studies are required to elucidate these relationships and their clinical relevance.

Using an explainable AI approach (SHAP), we established relationships between gene expression and predicted glioma subtype, as well as between expression and predicted OS (Figure 4, Supplementary Figures S2 and S3). The identified associations are consistent with the literature and indicate the correctness and biological interpretability of our models. Explainability is critical for AI systems designed for medical diagnostics. First, it ensures transparent logic behind AI predictions, which helps to strengthen the trust of patients and oncologists. Second, explaining prognoses allows clinicians to validate results against their existing clinical knowledge and experience. And third, explainability makes it easier to debug models in the case of prediction errors, pointing out the sources of errors, and enabling refinement of the model.

The developed explainable models contribute to understanding the unique molecular characteristics of gliomas. The prediction system has the potential to be implemented in clinical practice as an additional tool to routine analysis of imaging, histopathology and genomics data. It could help oncologists confirm the diagnosis, which would allow more accurate treatment planning and impact on patient survival.

Despite the high diagnostic potential of the developed models, our study has some limitations. First, the search for optimal hyperparameters of the ML models was confined to a limited range near the default settings. This implies that the resulting models might not be globally optimal with respect to their performance metrics. Second, although there are numerous feature selection algorithms, we utilized only two of them. In addition, when selecting a subset of features, we retained the top 1% most important ones, but adjusting this percentage upward or downward could potentially improve the results. Third, some annotations in the databases could be outdated due to modifications to the glioma classification system. Therefore, the developed model might need an update when the databases are reviewed according to the current WHO classification. Fourth, the classification of glioma subtypes was performed solely on the basis of their main histological types, without considering the grade. Addressing this gap is important and requires future studies. Fifth, the survival prediction model was constructed on the same 13 genes that were previously used to classify glioma subtypes. Given the considerable variation in survival rates between subtypes, future studies should focus on identifying subtype-specific survival markers. Furthermore, our survival model did not take into account clinical markers such as IDH mutation status and MGMT promoter methylation, which could potentially influence survival prediction. Sixth, the training dataset lacked ethnic diversity, which might limit the generalizability of the results to global populations. Finally, although we tested the models on independent datasets, including one from an external repository, additional large-scale datasets from multiple centers are necessary to further evaluate the robustness of our models.

## 5. Conclusions

In summary, we developed and tested two explainable ML models: one to classify three glioma subtypes and the other to predict survival probability. Both models make predictions based on the expression of 13 genes. The results of the quality assessment showed that the models have high classification and discrimination accuracy and are well calibrated. The application of the SHAP approach allowed us to determine the general contribution of the expression levels of selected genes on the outputs of both models, as well as producing explanations for individual predictions. The resulting classification of glioma subtypes and the evaluation of patient survival could be useful to select a personalized treatment strategy and to improve prognosis at bedside.

## Author Contributions

Conceptualization, O.V., V.T. and M.I.; formal analysis, O.V.; investigation, O.V. and V.T.; methodology, O.V. and M.K.; software, O.V.; supervision, A.T. and M.I.; validation, O.V.; visualization, O.V.; writing—original draft preparation, O.V.; writing—review and editing, O.V., V.T., M.K., A.T. and M.I. All authors have read and agreed to the published version of the manuscript.

## Funding

This work was supported by the Ministry of Economic Development of the Russian Federation (grant No 139-15-2025-004 dated 17.04.2025, agreement identifier 000000C313925P3×0002).

## Institutional Review Board Statement

Not applicable.

## Informed Consent Statement

Not applicable.

## Data Availability Statement

All the data used in the current study are publicly available in the CGGA (http://www.cgga.org.cn/) repository (accessed on 20 June 2022) and the TCGA (https://portal.gdc.cancer.gov/) repository (accessed on 2 August 2022).

## Conflicts of Interest

The authors declare no conflicts of interest.

## Notes

### Competing Interest Statement

The authors have declared no competing interest.

### Summary of Updates

Table 1 expanded; Figure 1E added; new Supplementary Table added; new Supplementary Figure added; additional gene enrichment analysis performed; sections Simple Summary, Institutional Review Board Statement and Informed Consent Statement added; Discussion section expanded; additional sources of literature added; text readability improved.

