## Supplementary material for "Explainable machine learning models for glioma subtype classification and survival prediction": Supplementary Figures.pdf

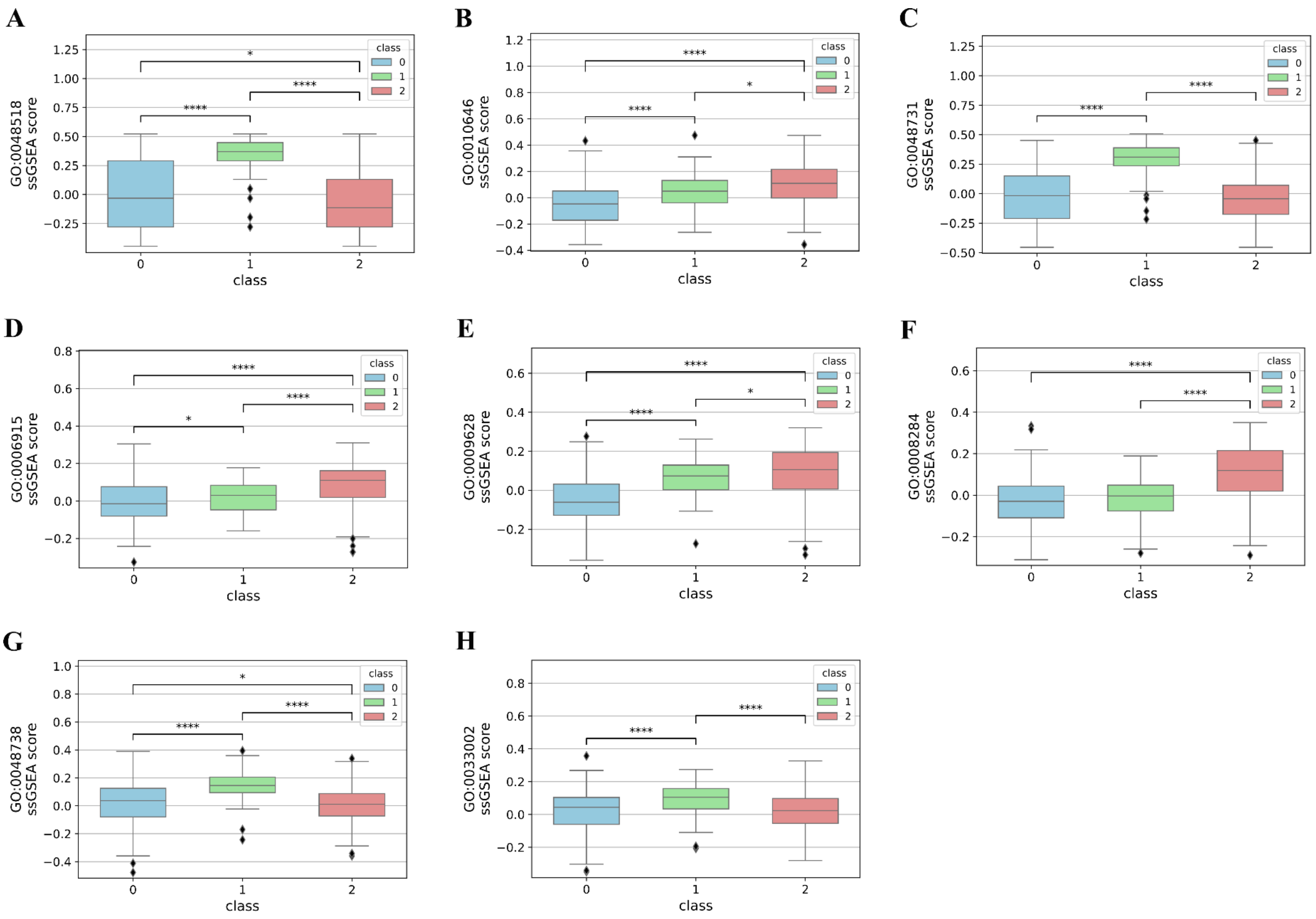


**Supplementary Figure S1.** Results of the single-sample gene set enrichment analysis (ssGSEA) for Gene Ontology (GO) terms including 13 selected genes. ssGSEA score for (**A**) GO:0048518, (**B**) GO:0010646, (**C**) GO:0048731, (**D**) GO:0006915, (**E**) GO:0009628, (**F**) GO:0008284, (**G**) GO:0048738, (**H**) GO:0033002. Class 0: astrocytoma, class 1: oligodendroglioma, class 2: glioblastoma. Post hoc pairwise Dunn's test: *p<0.05, **p<0.01, ***p<0.001, ****p<0.0001.

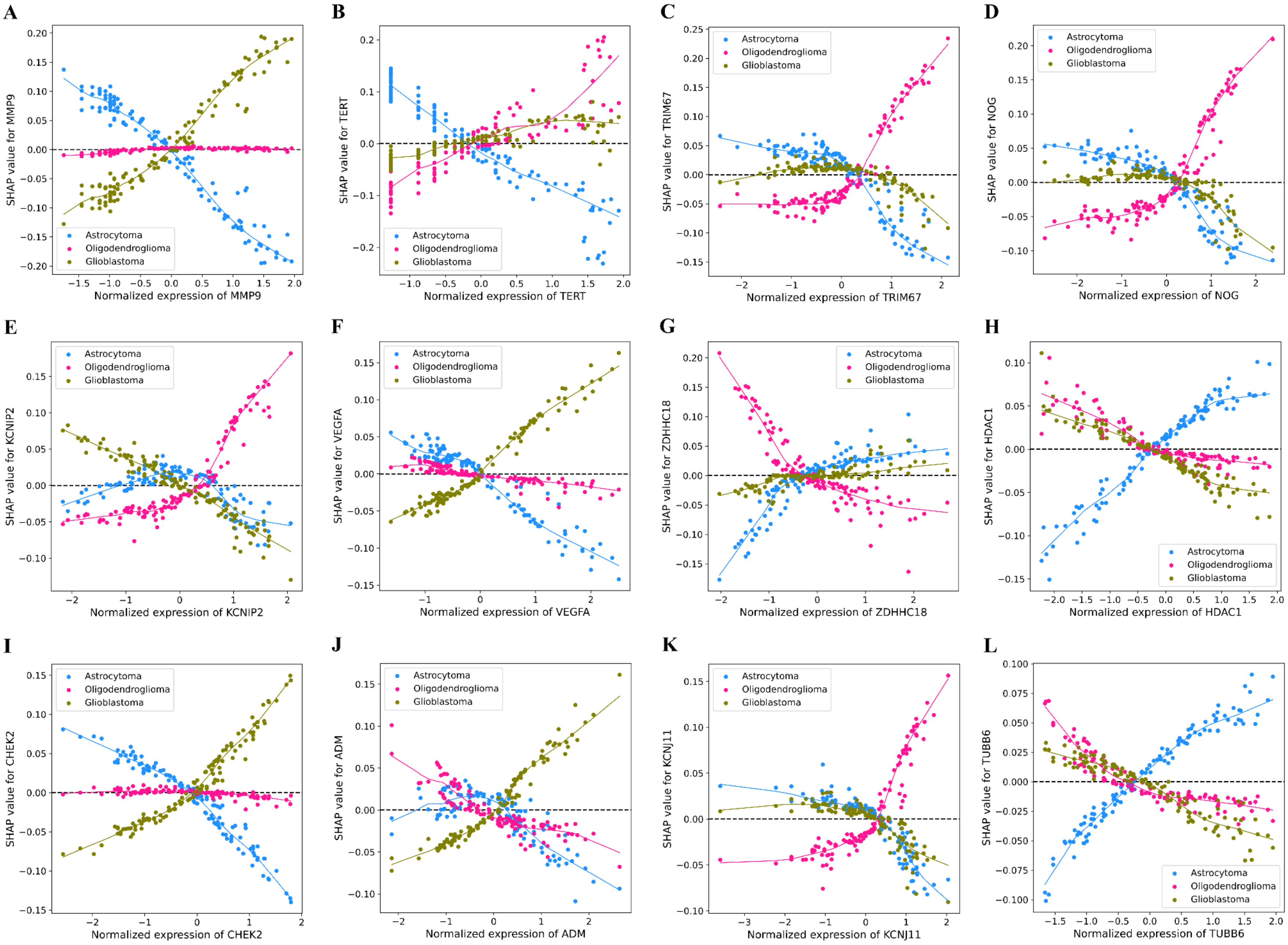


**Supplementary Figure S2.** Scatterplots between SHAP values and the expression levels of the genes in the classification model for three glioma subtypes: (**A**) MMP9, (**B**) TERT, (**C**) TRIM67, (**D**) NOG, (**E**) KCNIP2, (**F**) VEGFA, (**G**) ZDHHC18, (**H**) HDAC1, (**I**) CHEK2, (**J**) ADM, (**K**) KCNJ11, (**L**) TUBB6.

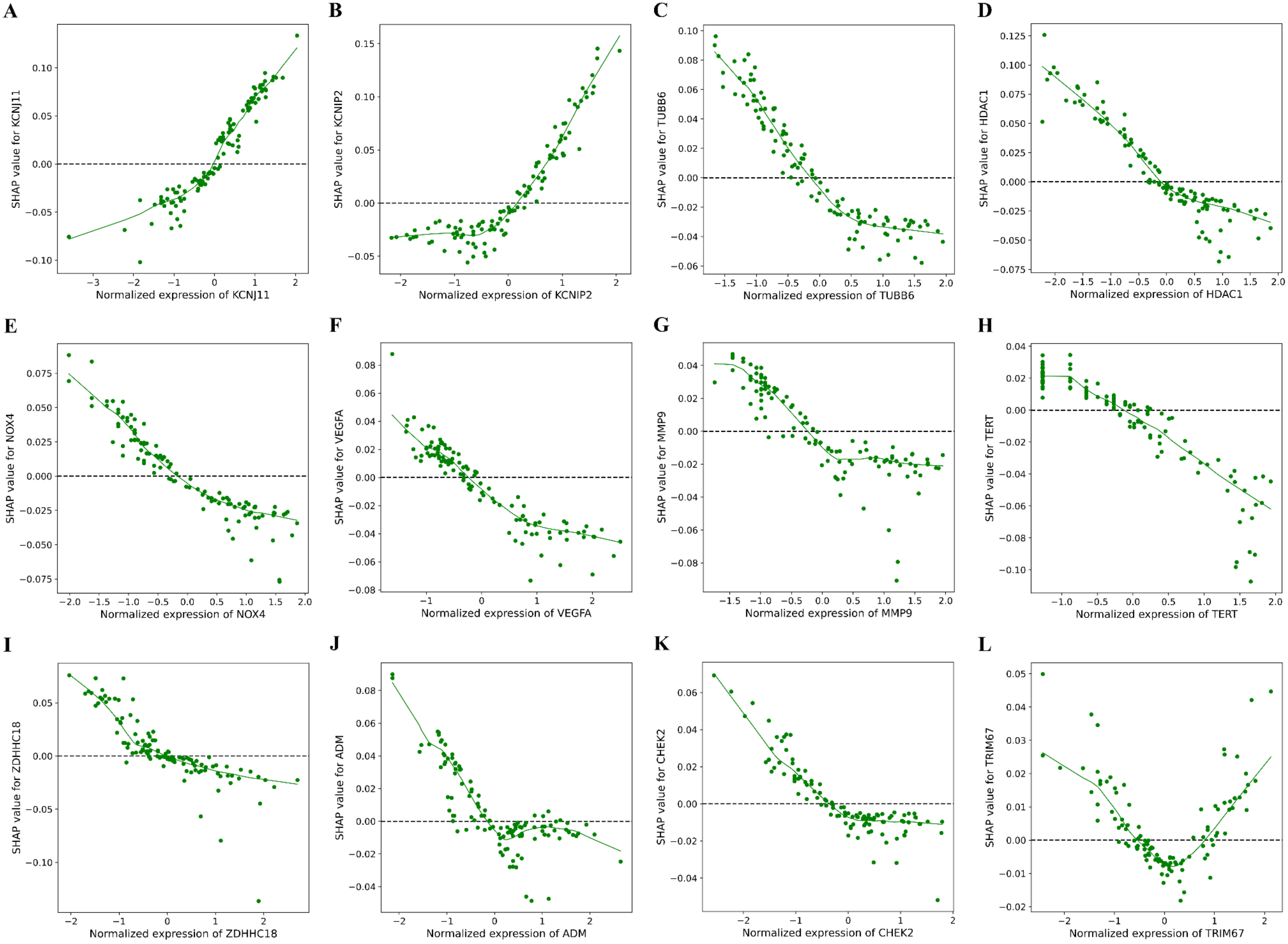


**Supplementary Figure S3.** Scatterplots between SHAP values and the expression levels of the genes in the model for predicting overall survival of patients with glioma: (**A**) KCNJ11, (**B**) KCNIP2, (**C**) TUBB6, (**D**) HDAC1, (**E**) NOX4, (**F**) VEGFA, (**G**) MMP9, (**H**) TERT, (**I**) ZDHHC18, (**J**) ADM, (**K**) CHEK2, (**L**) TRIM67.
